# Harnessing the adaptive potential of mechanoresponsive proteins to overwhelm pancreatic cancer dissemination and invasion

**DOI:** 10.1101/190553

**Authors:** Alexandra Surcel, Eric Schiffhauer, Dustin Thomas, Qingfeng Zhu, Kathleen DiNapoli, Maik Herbig, Oliver Otto, Jochen Guck, Elizabeth Jaffee, Pablo Iglesias, Robert Anders, Douglas Robinson

**Author notes:** Corresponding authors: Email addresses (Alexandra Surcel), (Douglas Robinson).

## Abstract

Metastatic disease is often characterized by altered cellular contractility and deformability, lending cells and groups of cells the flexibility to navigate through different microenvironments. This ability to change cell shape is driven in large part by the structural elements of the mechanobiome, which includes cytoskeletal proteins that sense and respond to mechanical stimuli. Here, we demonstrate that key mechanoresponsive proteins (those which accumulate in response to mechanical stress), specifically nonmuscle myosin IIA and IIC, α-actinin 4, and filamin B, are highly upregulated in pancreatic ductal adenocarcinoma cancer (PDAC) and in patient-derived pancreatic cancer cell lines. Their less responsive sister paralogs (myosin IIB, α-actinin 1, and filamin A) show a smaller dynamic range or disappear with PDAC progression. We demonstrate that these mechanoresponsive proteins directly impact cell mechanics using knock-down and overexpression cell lines. We further quantify the nonmuscle myosin II family members in patient-derived cell lines and identify a role for myosin IIC in the formation of transverse actin arcs in single cells and cortical actin belts in tissue spheroids. We harness the upregulation of myosin IIC and its impact of cytoskeletal architecture through the use of the mechanical modulator 4-hydroxyacetophenone (4-HAP), which increases myosin IIC assembly and stiffens cells. Here, 4-HAP decreases dissemination, induces cortical actin belts, and slows retrograde actin flow in spheroids. Finally, mice having undergone hemi-splenectomies with PDAC cells and then treated with 4-HAP have a reduction in liver metastases. Thus, increasing the activity of these mechanoresponsive proteins (in this case, by increasing myosin IIC assembly) to overwhelm the ability of cells to polarize and invade may be an effective strategy to improve the five-year survival rate of pancreatic cancer patients, currently hovering around 6%.

## Introduction

Altered mechanical states underlie morphological changes concomitant with cancer progression^1-7^ in two major ways. First, mechanical modifications often result from physical changes in the extracellular matrix (ECM) of the stroma and changes in the cellular composition of tumor microenvironments^8^. Second, the intrinsic genetic and proteomic compositions of cancer cells also impact their ability to navigate away from primary tumors, traverse mechanically disparate tissue layers, and establish metastatic niches. To respond to and eventually overcome physical and changing ECM barriers, migrating malignant cells (or collections of cells) must rely on their toolbox of cytoskeletal proteins. This toolbox that endows cells with their structural integrity, including their ability to sense and respond to their physical environment, along with their regulatory components, are collectively known as the mechanobiome. Unsurprisingly, this mechanical network undergoes striking changes in expression during cancer progression, which facilitates the dramatic spatial and temporal reorganization of the cytoskeleton intrinsic in metastasis. These altered expression patterns likely confer metastatic cells with the improved ability to deform, contract, and protrude into surrounding tissue.

Varying protein levels of critical components of the mechanobiome and the broader actin cytoskeleton have been observed in a wide range of cancers. For example, members of the formin and coronin families are upregulated, including in pancreatic cancer^9^, while cofilin overexpression is correlated with a poor prognosis among breast cancer patients^10^ and is upregulated in cervical^11^ and esophageal^12^ cancers. The overexpression of the actin crosslinking protein α-actinin 4 is likewise associated with poor patient outcomes in pancreatic^13^, colorectal^14,15^, gastric^16^, lung^17,18^, and breast^19^ cancers. Similarly, filamin B enhances the invasiveness of cancer cells into collagen matrices^20^. In addition, major cancer drivers and signaling proteins also have altered expression patterns and additionally impact cell mechanics. Yes-Associated Protein (YAP), whose overexpression is associated with numerous cancers^21^, modulates cellular actin architecture and nonmuscle myosin II regulatory light chain expression and phosphorylation, in turn affecting cortical tension and elasticity^22^. Early activating KRAS mutations that occur in over 90% of pancreatic cancers, as well as at high rates in colorectal and lung cancers, lead to increased deformability and altered contractility (*e.g*. ^23, 24^). Overexpression of members of the 14-3-3 family is negatively correlated with prognosis for liver^25^, pancreatic^26^, lung^27^, glioblastoma^28^, and other cancer patients. While 14-3-3 proteins are involved in numerous biological processes, they also modulate nonmuscle myosin II bipolar filament assembly and cell mechanics^29^ (West-Foyle *et al*., unpublished observations). Furthermore, a key inhibitor of myosin II, the myosin light chain phosphatase subunit MYPT1, is highly upregulated in pancreatic cancer^30^.

Collectively, the cell’s mechanobiome forms a mechanical continuum with the surrounding tissue and the relatively stiff nucleus to initiate and maintain metastatic motility. These proteins affect cell mechanics by impacting active force generation from actin assembly that pushes outward on the membrane, and myosin II contractility that pulls inward on the membrane. Myosin II contractility depends on other actin crosslinking proteins in the cytoskeletal network, and their cross-talk fine-tunes the deformability and contractility of the cell^31^. When alterations in the expression of these proteins occur, often due to key genetic lesions, the changes in mechanoresponsiveness lead to aberrant cell behavior^24^.

Here we test our hypothesis that expression patterns of key mechanoresponsive proteins and their sister paralogs change during PDAC progression, leading to altered deformability, contractility, and mechanoresponsiveness. We demonstrate that predicted mechanoresponsive proteins are upregulated in patient-derived pancreatic cancer tissue samples and cell lines, and that these proteins directly impact cell mechanics. We show that altered PDAC mechanics emanate in part from a changing ratio of nonmuscle myosin IIs, wherein myosin IIA and IIC are upregulated and myosin IIB is downregulated. We quantify the concentration of nonmuscle myosin paralogs in pancreatic cancer cells, and find that despite its relatively low concentration, myosin IIC has a significant impact on single cell behavior and collective behavior in tissue spheroids. We further probe the role of myosin IIC with a small molecule mechanical modulator, 4-hydroxyacetophenone (4-HAP), which increases the assembly of myosin IIC and stiffens PDAC cells^32^. We find that 4-HAP induces cortical actin belts and increases transverse actin arcs in single cells and tissue spheroids in a myosin IIC-dependent manner. This 4-HAP-induced change in cytoskeletal structure and mechanics leads to a decrease in PDAC metastasis in a mouse hemi-splenectomy model, demonstrating that specifically targeting elements of the mechanobiome, here by increasing their activity, has therapeutic potential for patients.

## Results

### Mechanoresponsive machinery is upregulated in pancreatic cancer

Using the social amoeba *Dictyostelium discoideum*, we previously identified ten proteins from a survey of ~35 proteins that accumulate in varying degrees to externally applied mechanical pressure^31,33,34^. Of these ten, we developed the physical theory accounting for the accumulation of three critical proteins – nonmuscle myosin II, α-actinin, and filamin – which are widely expressed structural elements of amoeboid and metazoan cells^31,35^. Our theory was able to predict which of the multiple paralogs of these protein families found in mammalian systems were mechanoresponsive^36^. To determine if these mechanoresponsive proteins behave similarly in pancreatic cancer cells, we assessed their localization in response to applied external stress via micropipette aspiration (MPA) (Fig. 1). Myosin IIA (MYh9), IIB (MyH10), and IIC (MYH14), as well as α-actinin 1 (ACTN1) and α-actinin 4 (ACTN4), were transiently expressed in several cell lines. These human-derived lines included HPDE (immortalized Human Pancreatic Ductal Epithelial cells), Panc10.05 (stage II pancreatic adenocarcinoma-derived), and AsPC-1 (stage IV ascites-metastasis-derived) (**Fig. 1A**). Cells were deformed for five minutes at a pressure of 0.3 nN/μm^2^, and the maximal protein accumulation in response to the dilational deformation at the aspirated tip of the cell was quantified by normalizing the fluorescence intensity at the tip region (I_p_) to the unstressed cortex opposite of the pipette (I_o_) (Fig. 1B).

**Figure 1:**
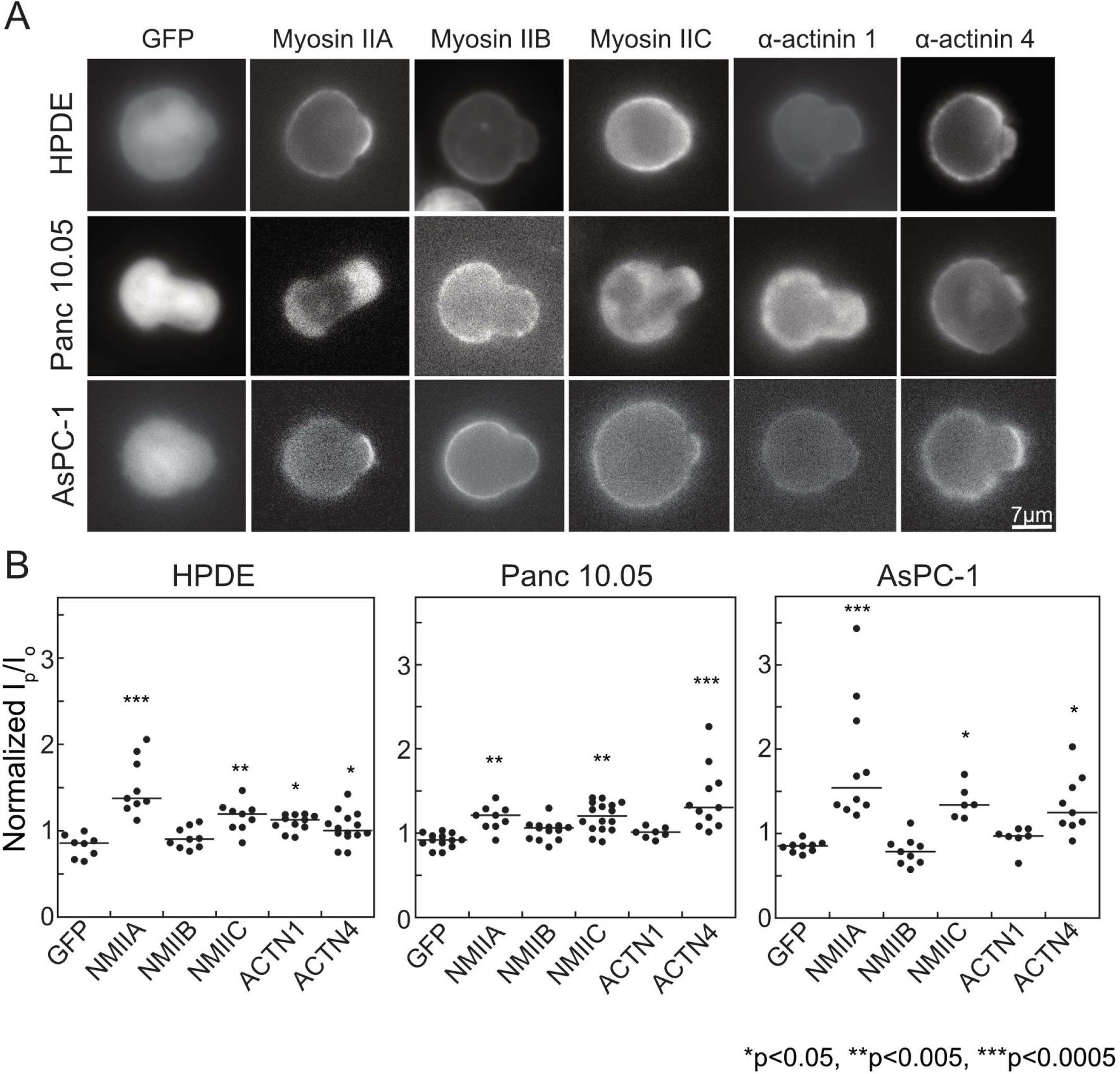
Nonmuscle myosin IIA, myosin IIC, and α-actinin 4 show mechanoresponsiveness in pancreatic cancer cells. **(A)** Representative images across HPDE, Panc10.05, and AsPC-1 cell lines of the maximum accumulation of GFP alone, GFP-labeled myosin IIs and α-actinin 4, and mCherry-α-actinin 1, show peak intensity after applied stress in MPA mechanoresponse experiments. Scale bar = 7 μm. **(B)** Quantification of mechanoresponsiveness normalized as a ratio of fluorescence intensity at the tip (I_p_) to the intensity at the opposite cortex (I_o_). Medians plotted; *p<0.05, **p<0.005, ***p<0.0005.

In all cell lines, compared to the GFP-vector control, myosin IIA and myosin IIC were mechanoresponsive, whereas myosin IIB showed no accumulation, consistent with its previously observed cell-type-specific mechanoresponsiveness^36^. Of the α-actinins, α-actinin 4, but not α-actinin 1, was mechanoresponsive, especially in Panc10.05 and AsPC-1 cells. This differential behavior between the α-actinin paralogs likely results from the much lower actin binding affinity of α-actinin 4 (K_d_=32 μM) compared to α-actinin 1 (K_d_=0.36 μM), which leads to more dynamic α-actinin 4 behavior necessary for the protein to respond to mechanical stress^36-37^. Likely due to their large size (~280 kD), transfection efficiency of filamin A and filamin B constructs was insufficient in all three of these pancreas-derived cell lines to assess mechanoresponsiveness.

We hypothesized that concomitant with cancer initiation and progression, the mechanoresponsive machinery would be upregulated to endow cells or collections of cells with the ability to sense and respond to changing physical environments across discrete tissue types. To test this idea, we performed immunohistochemistry on pancreatic cancer tissue samples from 20 patients across all seven proteins – the three nonmuscle myosin IIs (IIA, IIB, and IIC), the two α-actinins (1 and 4), and the two filamins (A and B). We compared normal ducts with cancerous ducts and metastatic lesions. In addition, we derived a scoring system that allowed us to delineate between high expression and low expression, as well as percentage of cells positively stained within the quantified ducts (outlined in **Sup. Fig. 1A**). All mechanoresponsive proteins showed a significant shift and increase in expression in cancerous versus normal ducts (Fig. 2, **Sup. Fig. 1B**). Myosin IIA and myosin IIC increased expression, with myosin IIC specifically upregulated in the adenocarcinoma, while myosin IIA increased across the pancreatic cancer stroma in addition to the ducts. The non-mechanoresponsive myosin IIB showed no significant change in expression, with very little staining in general. The mechanoresponsive α-actinin 4 also increased in expression concurrent with cancer progression, while α-actinin 1 maintained mostly uniform expression levels across ducts. Filamin B, which is mechanoresponsive in other cell lines^36^, is upregulated specifically in cancerous ducts. Filamin A is upregulated across the entire pancreatic tissue. These patterns are also noted in non-invasive lesions, termed pancreatic intraepithelial neoplasia (PanINs) (**Sup. Fig. 1B**), with increasing expression associated with cancer progression. Our results are largely in keeping with normal versus pancreatic cancer tissue datasets in the Gene Expression Omnibus (GEO) (**Sup. Fig. 1C**)^38^. The staining patterns are also consistent with the Human Protein Atlas which tracks RNA and immunohistochemistry and which suggests that filamin B and α-actinin 4 are poor prognostic indicators for pancreatic cancer patients^39^. Filamin A shows variable PDAC expression across these studies, and α-actinin 1 displays high staining in both normal and cancerous cells. Overall, the immunohistochemistry data indicate that, as a unit, the mechanoresponsive machinery is upregulated in the pancreatic ductal adenocarcinoma of patients.

**Figure 2:**
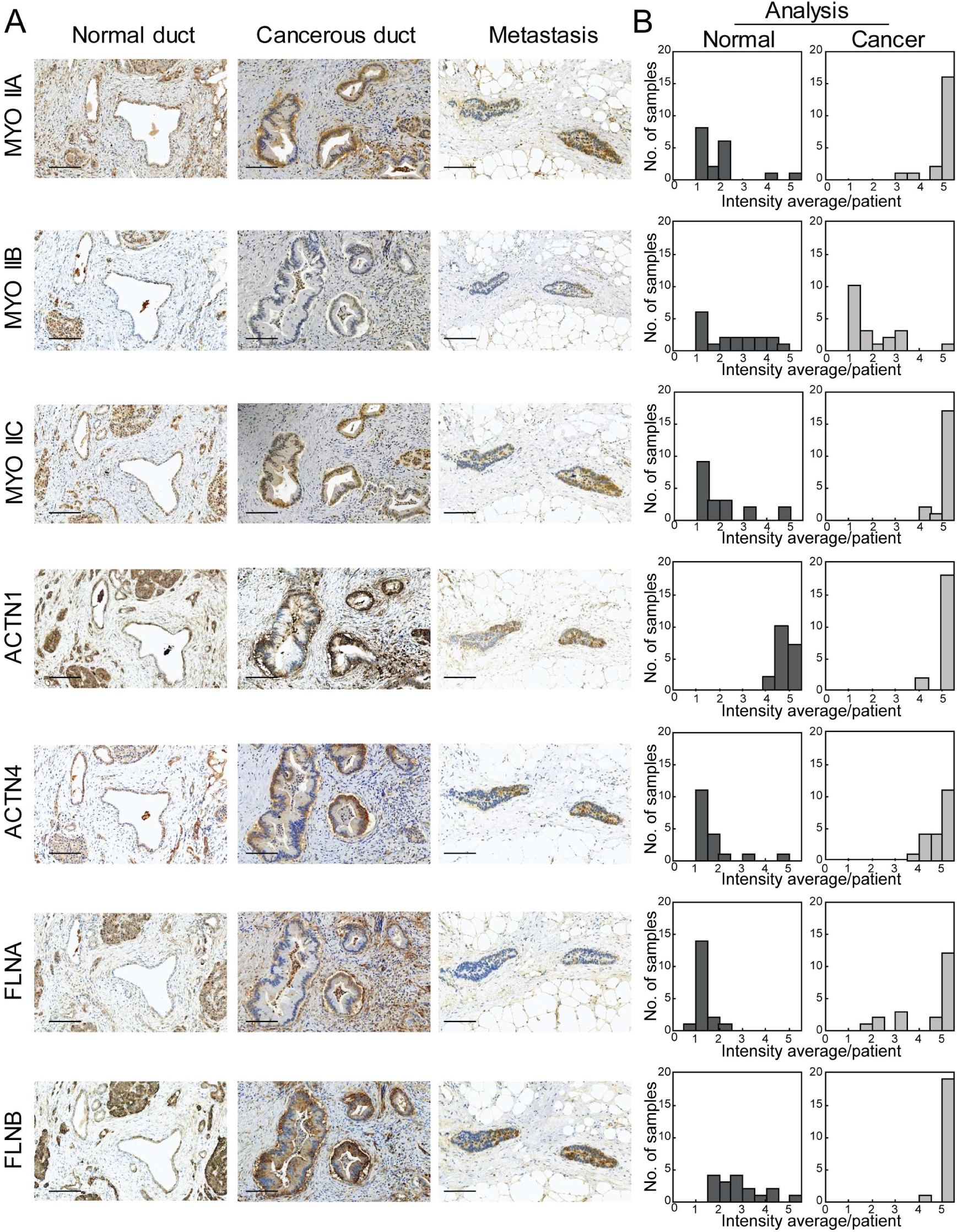
The mechanoresponsive machinery is elevated in pancreatic ductal adenocarcinoma in human pancreatic tissue. **(A)** Immunohistochemistry staining of pancreatic tissue from PDAC patients shows an increase in mechanoresponsive proteins nonmuscle myosin IIA and IIC, α-actinin 4, and both filamin A and B. Scale bar = 100 μm. For each sample – normal duct, cancerous duct, and metastatic lesion – the same site is shown across all seven antibodies stained. In addition, both the normal and cancerous ducts are from the same patient. **(B)** Quantification of staining intensity and surface area across 20 patients illustrate dynamic up-regulation and shift of mechanoresponsive proteins, as well as filamin A. Data is plotted as a histogram of the intensity average per patient (described in **Sup. Fig. 1A** and **Materials and Methods**) across the study group.

### PDAC cell lines can be used as a mechanoresponsive model for PDAC

To determine if PDAC cell lines can be used to study the changing mechanobiome landscape, we first assessed if the expression patterns that we observed in patient samples (Fig. 2) matched generally with changes between WT-like HPDE and various tumor and metastatically-derived lines. Western analysis across four lines revealed a general increase in myosin IIA and IIC, the disappearance of myosin IIB, and an increase in α-actinin 4 and filamin B, with moderate or unchanged levels of α-actinin 1 and filamin A (Fig. 3). Because the small molecule 4-HAP that we will use to modulate cell mechanics works through myosin IIB and IIC (see below), we measured the concentration of each myosin paralog in these pancreatic cancer-derived cells. We first calibrated both HeLa cells, which express myosin IIA and IIB, and AsPC-1 cells, which express myosin IIC, to generate a quantitative comparator for measuring each paralog’s concentration across cell lines. To calibrate HeLa and AsPC-1 cells, we added purified paralog-specific myosin II tail fragments to the extract (**Sup. Fig. 2A**). From these calibration measurements, we calculated that nonmuscle myosin IIA concentration in human pancreatic cells ranged from 540 nM in HPDE cells to 770 nM in Panc4.03 cells. These values compare favorably with the amounts of myosin II in budding yeast (Myo2p, 450 nM; Myp2p, 380 nM,^40^) and in *Dictyostelium discoideum* (Myo II, 3.4 μM,^41^) (Fig. 3B). By comparison to myosin IIA, myosin IIB and IIC are found at much lower concentrations. Interestingly, myosin IIC increased 1.5-fold from approximately two percent of all myosin II in HPDE cells to about three percent of all myosin II in AsPC-1 cells, while myosin IIB decreased from ~8% of all myosin in HPDEs to undetectable in AsPC-1 cells (Fig 3A,B). It is important to note that the HPDE cell line was immortalized by E6/E7 transformation, which affects p53 activity among other alterations, so the levels of the myosins in HPDEs may be changed compared to the original sample taken from the patient. The dramatic change in myosin IIC in normal and cancerous ducts in the immunohistochemistry supports the myosin quantification across PDAC cell lines (Fig. 2). While such small amounts of myosin IIC might suggest that the protein cannot impact cell mechanics and behavior significantly, the studies described below demonstrate otherwise.

**Figure 3:**
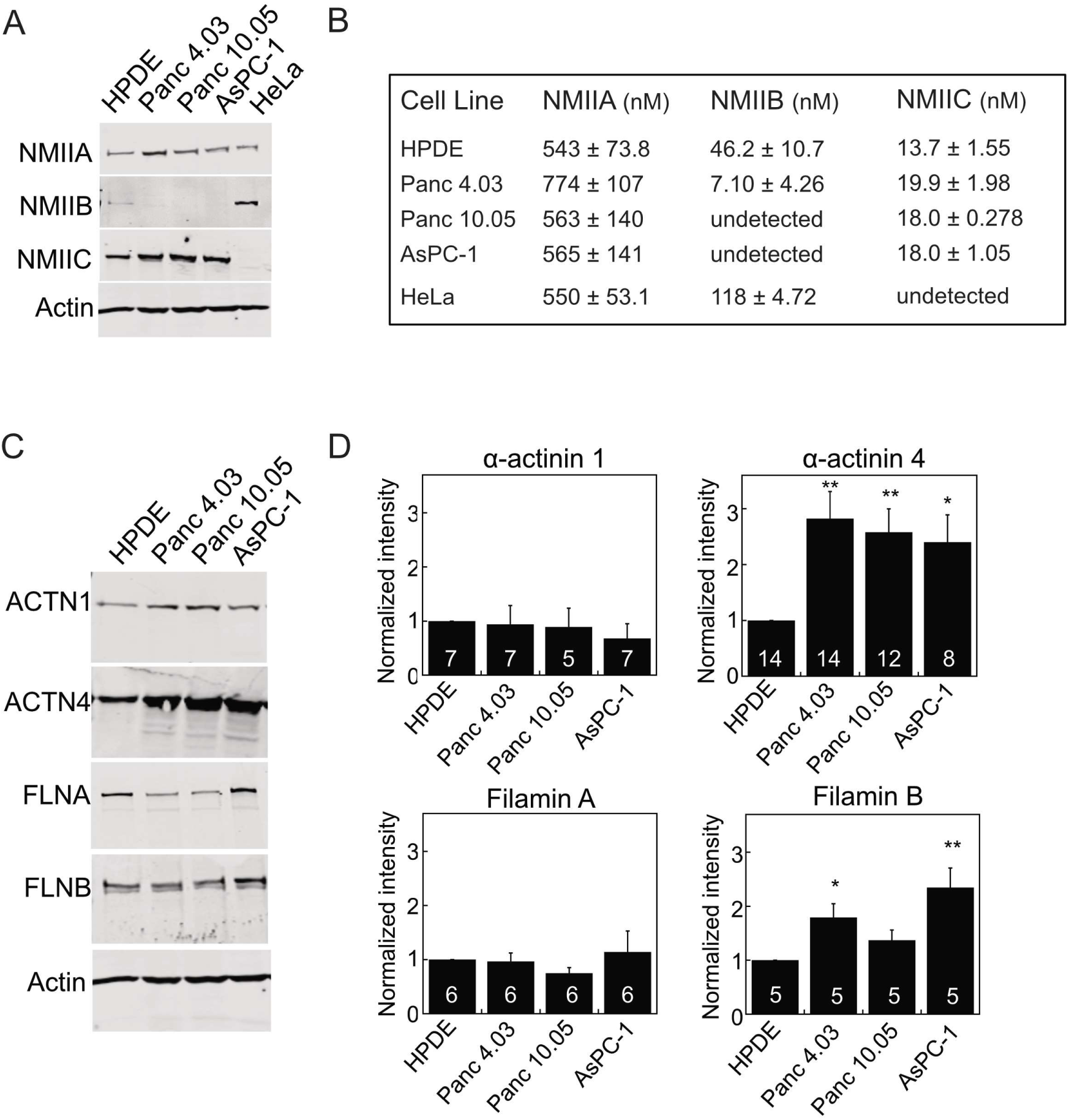
Mechanoresponsive proteins increase in pancreatic cancer-derived cell lines. **(A)** Expression of myosin IIA (NMIIA), myosin IIB (NMIIB), myosin IIC (NMIIC), and actin in HPDE (normal pancreatic ductal epithelium), Panc4.03 (stage II primary tumor), Panc10.05 (stage II primary tumor), and AsPC-1 (stage IV ascites metastasis) cells, compared with HeLa lysate for the purposes of quantification. NMIIA and NMIIC increase in expression, while NMIIB decreases in expression in cancer cell lines. **(B)** Quantification of the cellular concentration of each myosin II paralog. **(C)** Expression of alpha-actinin 4 (ACTN4) and filamin B (FLNB) increase, while expression of alpha-actinin 1 (ACTN1) and filamin A (FLNA) do not change in cancer cell lines. **(D)** Quantification of western blots, examples shown in **(C)**, where numbers on the bars indicate n-values. *p<0.05, **p<0.005.

### Mechanoresponsive machinery impacts cell mechanics in PDAC lines and can be modulated by 4-hydoxyacetophenone (4-HAP)

We previously demonstrated that WT-like HPDE cells are less deformable than patient-derived PDAC cell lines^32^. To determine if a subset of the mechanoresponsive elements of the PDAC mechanobiome contribute to this mechanical differential, we used micropipette aspiration (MPA) to measure the effective cortical tension (T_eff_) of cells with overexpression or knockdown of myosin II and α-actinin paralogs in Panc10.05 cells (Fig. 4A). Across all cell lines, knockdowns were 70-95% (**Sup. Fig. 2B**). Driving myosin IIA levels up or down yielded an altered cortical tension which rose and fell with myosin IIA expression levels. Myosin IIB overexpression had no impact on cortical tension. Myosin IIB knockdowns were not pursued since Panc10.05 cells (as well as other PDAC lines and patient tissue samples) have no detectable levels of this protein (Fig. 3A, Fig. 2A, **Sup. Fig. 1**). Alpha-actinin 1 overexpression had no effect on cell mechanics, while both overexpression and knockdown of α-actinin 4 decreased the T_eff_ by half (Fig. 4A). Interestingly, despite contributing only 3% of the overall myosin II in these cells (Fig. 3C), overexpression and knockdown of myosin IIC had a profound impact on cell mechanics, leading to an overall softening of the cellular cortex and an ~50% reduction in cortical tension. To further explore the impact that myosin IIC has on the PDAC mechanobiome and cell mechanics, we used the small molecule 4-hydroxyacetophenone (4-HAP) which we previously identified in a *Dictyostelium* screen for mechanical modulators. In *Dictyostelium*, 4-HAP increased cortical tension by driving myosin II to the cell cortex. 4-HAP shows myosin II paralog specificity in mammalian cells by increasing the assembly of myosin IIC (and IIB) and the elasticity of several PDAC cell lines^32^. As predicted, 4-HAP treatment increased the cortical tension of control cells where myosin IIC is present, but had no impact on myosin IIC-depleted cells (Fig. 4A).

**Figure 4:**
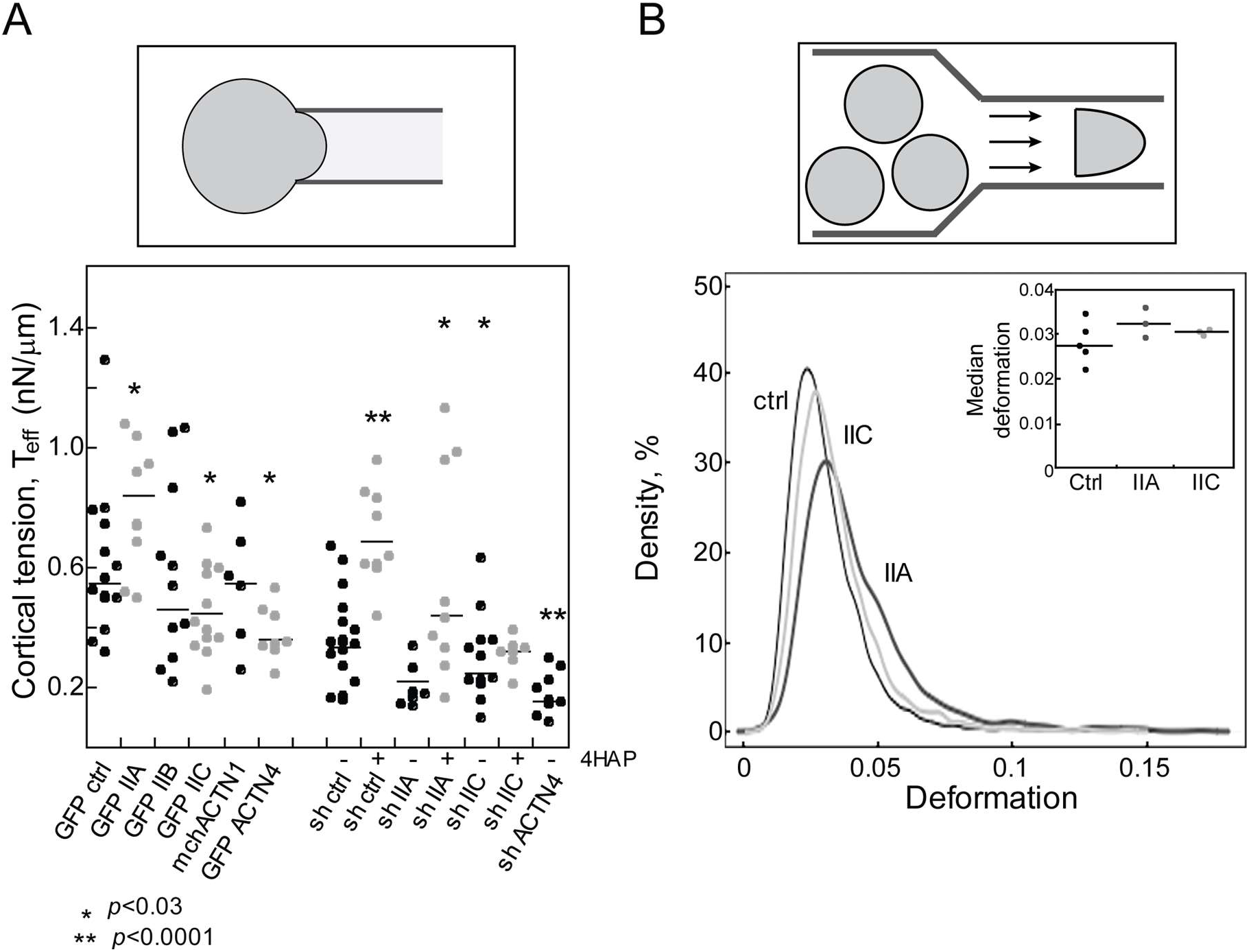
Mechanoresponsive proteins impact cortical tension and deformability of PDAC cells. **(A)** The effective cortical tension (Teff) measured by MPA experiments (schematic) is significantly affected by overexpression of myosin IIA, myosin IIC, and α-actinin 4, all mechanoresponsive paralogs. 500-nM 4-HAP increases the cortical tension of sh-control cells. 4-HAP also increases the cortical tension in myosin IIA knockdown cells, whose primary myosin paralog myosin IIC is activated by 4-HAP ^32^. Knockdown of myosin IIC also leads to a decrease in cortical tension, unchanged by 4-HAP challenge. Medians plotted; * p<0.03; **p<0.0001 relative to control. (B) RT-DC experiments (schematic) demonstrate increased deformation when myosin IIA and myosin IIC are knocked-down (plotted as a probability distribution). All cell lines are generated from Panc10.05 cells. N=7521 cells for control, 1383 cells for myosin IIA knockdown, and 5933 cells for myosin IIC knockdown. Cell types are distinct (p<0.0001). Inset: Median deformation of three to five RT-DC independent runs across cell types from which the elastic modulus was calculated.

In addition to micropipette aspiration, which measures mechanical properties on the >500-ms time-scale (cortical tension measurements are performed over 10s of seconds), we used Real Time Deformability Cytometry (RT-DC)^42,43^, which measures mechanics on the 4-ms timescale and across the whole cell (**Sup. Fig. 3**). Reduction of myosin IIA, but not myosin IIC, increased cell deformation and altered elasticity (control, 1.2 kPa, shIIA, 1.1 kPa, and shIIC 1.2 kPa; Fig 4B). The small 8% reduction in the elastic modulus measured for knockdown of myosin IIA, which is the most abundant paralog in the PDAC cells, is similar to the reduction in elasticity measured for WT versus *myoII* genetic deletion cells in *Dictyostelium*^44^.

### Nonmuscle myosin IIC alters actin bundling in cells, impacting migration and dissemination

Nonmuscle myosin II proteins modulate cell-cell and cell-substrate adhesions, tune the mechanical profile of cells by affecting such parameters as cortical tension and elasticity, and impact cytoskeletal arrangements. To determine how myosin IIC specifically impacts cytoskeletal organization in collectives of cells, particularly as the minor contributor to total myosin II in PDAC, we generated tissue spheroids with knockdown Panc10.05 cell lines. Control knockdowns plated in 3D collagen-I matrices showed partial dissemination, which was greatly increased in NMIIA knockdowns, similar to previous observations^45-47^. Knockdown of myosin IIC showed no dissemination (Fig. 5A). When myosin IIC was activated in spheroids upon treatment with 4-HAP for 24 hours, dissemination in both the control and NMIIA knockdown reduced, while no effect was observed in the myosin IIC depleted spheroids. To better visualize structure, spheroids were plated on collagen-I coated 2D substrates and similar morphological differences and changes were also observed as in the 3D cultures (Fig. 5A, row 2). In addition, we observed tight actin cortical banding patterns on the periphery of the tumor spheroids in the absence of myosin IIC. This banding pattern was also observed in the control and NMIIA knockdown spheroids with 4-HAP treatment. To assess this actin redistribution, we used computer-assisted image analysis on the 2D spheroids and measured the continuity of banding, the percentage coverage of actin filaments at the spheroid edge, and the homogeneity of the actin in those structures (calculated as the standard deviation of pixel intensity) (**Sup. Fig. 4**). Across all analyses, myosin IIA knockdown spheroids had the least amount of discrete and continuous banding (Fig. 5B, **Sup. Fig. 4B**) and the least amount of actin staining at the tissue edge (**Sup. Fig. 4C**). Upon 4-HAP treatment, all of these metrics changed – more discrete and continuous belts emerged and the median percentage staining of actin increased 2-fold. By comparison, the myosin IIC knockdown tissue spheroids had additional discrete and continuous actin belts that remained unchanged upon 4-HAP addition (Fig. 5B, **Sup. Fig. 4B,C**), consistent with 4-HAP working primarily through myosin IIC. Interestingly, by these analytics, treatment with 4-HAP in the control tissue spheroids seemed to decrease discrete band formation.

**Figure 5:**
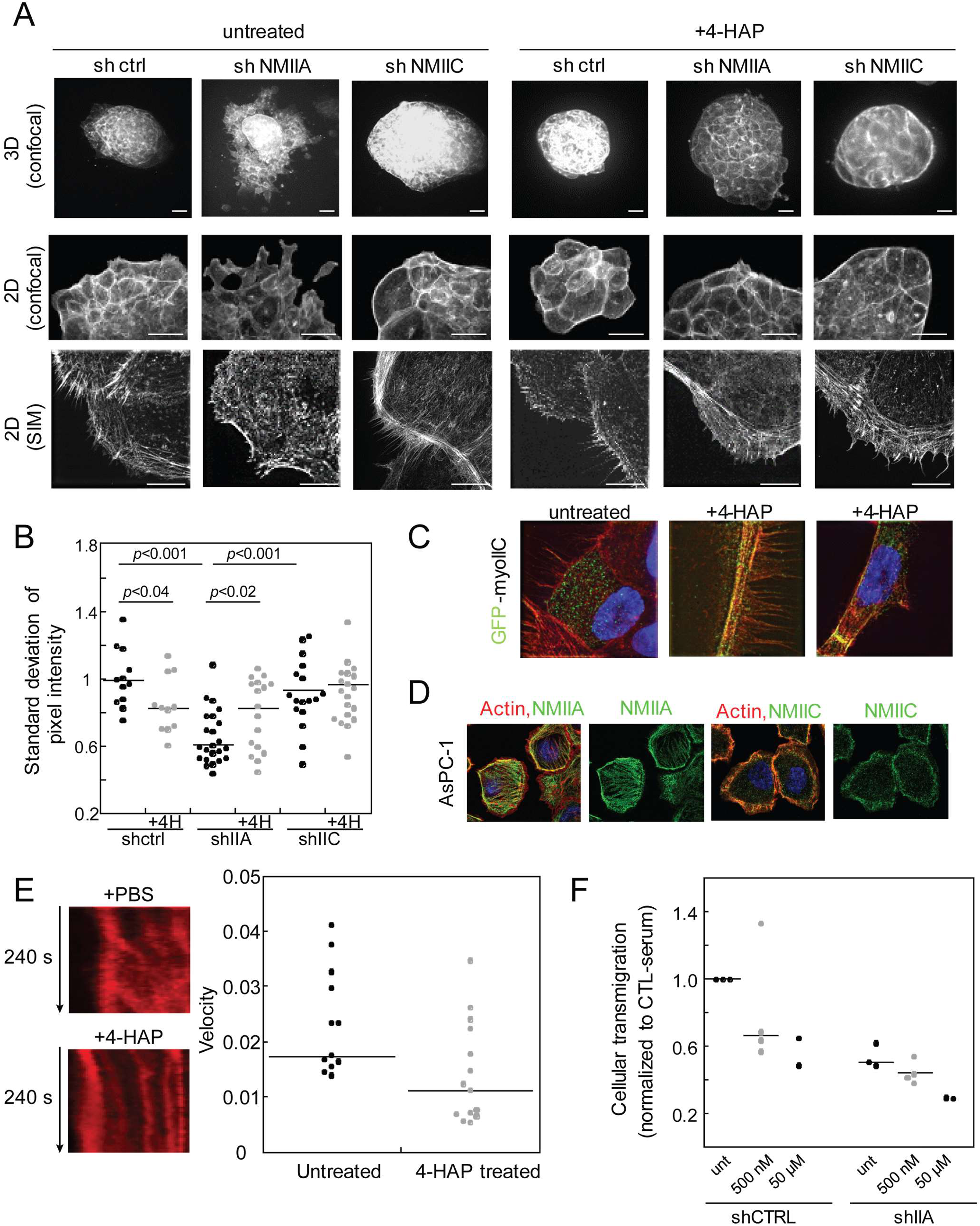
Myosin IIC impacts cytoskeletal actin architecture, leading to actin belt formation and altered cell behavior upon 4-HAP treatment. **(A)** Tissue spheroids of Panc10.05 derived cells, grown and imaged in 3D (Matrigel) or 2D (collagen) show dissemination, exacerbated in myosin IIA knockdowns, and reversed in the absence of myosin IIC, with the emergence of actin cortical belts. Actin belts are formed in 4-HAP treatment when myosin IIC is present. Scale bar = 40 μm except SIM images in which case scale bar = 10 μm. **(B)** Quantification and verification of these actin structures as a standard deviation of pixel intensity (a coarseness index); see Methods section for detailed explanation. Medians plotted; significance on graphs. **(C)** GFP-myosin IIC decorates actin filaments, especially actin belts generated by 4-HAP treatment in tissue spheroids. **(D)** Endogenous labeling of myosin IIA in fixed AsPC-1 cells shows myosin IIA colocalized on actin stress fibers and myosin IIC colocalized with actin on the cell cortex. **(E)** Sample kymographs of line scans across active leading edges in AsPC-1 SirAct live-stained cells. Graph shows decreased retrograde flow with 4-HAP treatment. Medians are plotted on the graph. **(F)** Treatment with 4-HAP of AsPC-1 shCTRL and shIIA cells shows moderate dose-dependence reduction of transwell migration. Medians are plotted on the graph.

We then used Structural Illumination Microscopy (SIM) to acquire higher resolution views of the structures along the tissue spheroid edges (Fig. 5A **bottom row, B**). In the control spheroids, the actin belts are formed by the structural rearrangement of the actin bundles. Here, 4-HAP induces a coarser distribution of actin composed of dense actin belts and leads to the emergence of filopodial-like structures or retraction fibers. In the myosin IIA knockdowns treated with 4-HAP, tight actin belts, as well as elaborate arrays of parallel actin bundles, are also clearly visible. In contrast, myosin IIC knockdowns had peripheral actin structures that were largely unchanged between untreated and 4-HAP-treated samples. Thus, 4-HAP induces alterations in peripheral actin structures in a myosin IIC-dependent manner.

To address myosin IIC’s role in actin rearrangements, we next determined its cellular localization. In tissue spheroids, fluorescently-labeled myosin IIC is both diffusely localized throughout the cell and along actin filaments (Fig. 5C). When actin filaments collapse to form actin belts upon 4-HAP treatment, myosin IIC shows strong co-localization with those belts. In single Panc10.05 and AsPC-1 cells, endogenous myosin IIC is predominately confined to the cell cortex including in actin-rich protrusions, whereas myosin IIA localizes along stress fibers (Fig. 5D).

Overall, these observations and analysis highlight two major findings. First, in collections of cells, despite being present in small quantities, myosin IIC plays a role in actin rearrangements and dynamics. Myosin IIA, known to contribute to retrograde actin flow, may work in concert with myosin IIC to drive these cytoskeletal rearrangements. Second, 4-HAP reduces the fluidity of the network: by stabilizing actin belt structures in the absence of myosin IIA and in the presence of both myosin IIA and myosin IIC, 4-HAP alters the actin cytoskeleton in a manner that inhibits dissemination from tissue spheroids.

Because retrograde flow moves actin away the cell perimeter, the cortical actin belts could be formed by a reduction in this flow. Therefore, we examined the impact of 4-HAP on retrograde flow. We live-stained cells with SiR-actin and observed flow using lattice sheet microscopy. Most 4-HAP-treated cells had undetectable levels of actin flow. Those in which retrograde flow could be measured showed 50% reduction in velocity over untreated controls (Fig 5E, **Sup. Fig. 5, Sup. Movies 1, 2**). This large and measurable impact on actin dynamics explains in part the dose-dependent effect on trans-well migration in control and myosin IIA knockdown AsPC-1 cells (Fig 5F). Overall, 4-HAP’s impact on actin flow, cell dissemination, and invasion suggests its potential for reducing PDAC metastasis.

### 4-HAP decreases PDAC metastatic potential in mouse model

Critical regulators of myosin II have significantly altered expression associated with pancreatic cancer progression and in pancreatic cancer-derived cell lines^9,29,30^. These genetic alterations suggest that the activation of myosin II, particularly myosin IIC whose expression is specifically elevated in pancreatic ductal epithelia, may be an attractive target for impacting PDAC cell behavior. Therefore, we tested 4-HAP in a mouse model for PDAC metastasis. Hemi-splenectomies with 4-HAP pre-treated metastatic AsPC-1 cells were performed on *nude* mice which were subsequently divided into three groups: control, PBS (200 μl PBS IP injections, every other day), and 4-HAP (5 mg/ml, 200 μl IP injections, every other day). Mice were harvested at 5-weeks post-surgery, when the first mouse expired. Metastasis to the liver was observed and quantified by an in-house Matlab script that determined surface area tumor coverage. Results from both liver weights and tumor coverage show that 4-HAP treatment leads to a reduction in tumor formation (Fig. 6, **Sup. Fig. 6**).

**Figure 6:**
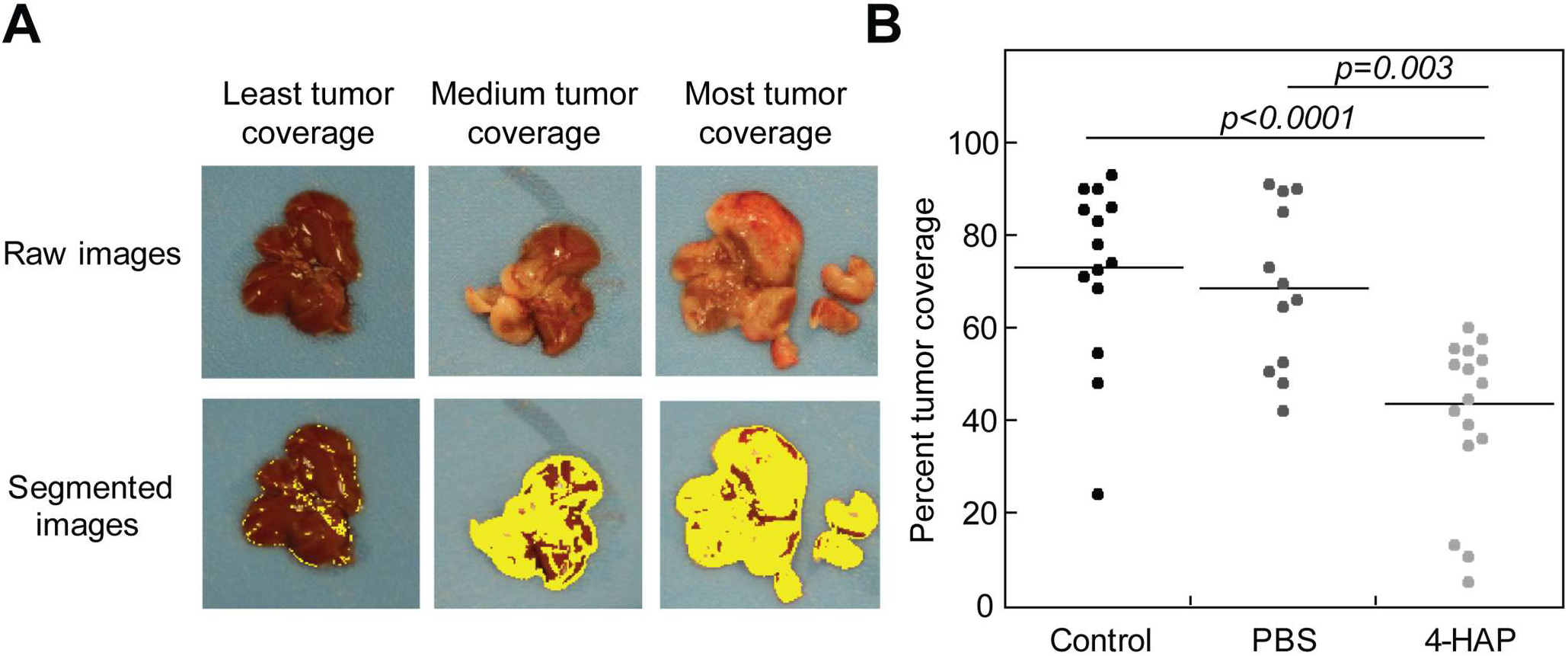
4-HAP reduces PDAC liver metastasis in murine model. Livers harvested from 4-HAP-treated mice that underwent hemi-splenectomies with AsPC-1 cells show a reduction in tumor coverage over those livers from untreated mice. **(A)** Images were quantified using image segmentation based on color gradients to discern tumor coverage (**Sup. Fig. 6**). **(B)** Quantification of tumor coverage show a 50% reduction in surface tumor load. Medians are provided on the graph.

## Discussion

A cell’s ability to react to changing mechanical and chemical cues in its environment depends on the adaptability of its mechanobiome. Increased contractility and altered deformability, as well as rapid turnover of cytoskeletal proteins, are trademarks of cells responding to constantly changing surroundings. Entire programs are upregulated to provide cells with added adaptability at specific time points as developing embryos and differentiating cells show increased expression of mechanoresponsive proteins during mechanically turbulent periods. For example, filamin’s mechanoresponsiveness is required for maturation of actin-rich ring canals that interconnect the nurse cells and oocyte in developing *Drosophila* egg chambers^48^, and filamin B is upregulated in embryonic vascular endothelial cells^49^. Both α-actinin 1 and α-actinin 4 show temporally defined expression in developing zebrafish embryos, with both expressed in the notochord and α-actinin 4 also expressed in the developing gut^50^. Each of the nonmuscle myosin II paralogs have roles in development as well, including but not limited to neurite outgrowth and maturation (NMIIB and NMIIC)^51^, nephron development (NMIIA and NMIIB) (*e.g*.^52^), and hearing (NMIIA)^53^.

In the mechanobiome, forces are shared between myosin II and different actin crosslinkers, with myosin II having potentiating or inhibitory effects on certain crosslinkers and vice versa^31,54^. This mechanosensory system constitutes a control system where mechanical inputs can be converted to signaling outputs in a manner analogous to chemical signal transduction^34,55^. Through our work^31^, an important delineation has emerged: the cell has at least two systems of proteins that when depleted, lead to a reduction in cortical viscoelasticity and tension. One set of proteins leads to increased mechanoresponsiveness, while the other set of proteins leads to reduced mechanoresponsiveness. Cells that mature into terminally differentiated tissues often readjust the cytoskeletal milieu to favor reduced mechanoresponsiveness over their developmental program. These stable expression patterns, however, are altered in precancerous cells (often caused by upstream genetic lesions), and revert cells to programs activated in early development that endow them again with increased adaptability.

Here we show that in the case of pancreatic ductal adenocarcinoma, the mechanoresponsive proteins myosin IIA, myosin IIC, α-actinin 4, and filamin B are upregulated in patient-derived tissues; they alter the structural arrangement of the actin cytoskeleton and impact cell mechanics. In addition, despite its low abundance, myosin IIC works in conjunction with myosin IIA to facilitate actin organization and retrograde flow. These data suggest that the interplay between these two paralogs is necessary to control leading edge dynamics of PDAC cells, bestowing a mechanical advantage to these cells. Also, our data on myosin IIC suggests that changes in the expression level of minor proteins that are often discounted in larger data mining may in fact, be worthy of reconsideration, given the large impact myosin IIC has on cell mechanics and cell behavior. Our *in vivo* metastasis assays demonstrate that the myosin IIA-IIC dynamic can be fine-tuned towards a therapeutic benefit with mechanical modulators such as 4-HAP. In addition, because myosin IIC is specifically upregulated in ductal adenocarcinoma cells, the pharmacological modulation of this protein is unlikely to negatively impact healthy pancreatic tissue; 4-HAP should synergize with other strategies such as immunological intervention in pancreatic cancer patients, since immune cells do not typically express myosin IIC.

Furthermore, the observations presented here imply that targeting cancer by broadening strategies to include small molecule activators instead of mainly inhibitors, can have significant effects on metastatic load, and ultimately patient survival. Promoting additional activation of mechanoresponsive proteins has several advantages. First, we can fine-tune the activation of proteins that are upregulated by cancerous tissue, thus harnessing the cell’s intrinsic protein make-up to revert them to more normal phenotypes, while protecting healthy cells that do not harbor upregulation of targeted proteins. Second, this strategy draws upon the normal biochemistry of the protein to overwhelm the system. In our studies, we are hitting on both points: We are using increased the activation (via overassembly) of myosin IIC, which is specifically upregulated in the PDAC, to overcome the protein’s innate adaptive ability. 4-HAP treatment converts myosin IIC into a stabilizer that inhibits the tumor cell’s ability to polarize and reorganize its actin cytoskeleton. Overall, incorporating the mechanobiome as a targetable drug space in combination with other therapeutic approaches, is likely a viable strategy for reducing PDAC metastasis.

## Materials and Methods

### Cell culture and strains

#### Parental pancreatic cell lines

Human pancreatic ductal epithelial cells (HPDE) were obtained from Dr. Ming-Sound Tsao (University of Toronto, Ontario Canada) and human primary tumor-derived cells (Panc10.05) and human metastatically-derived cells (AsPC-1) were purchased from ATCC. Panc02 cells (highly tumorigenic murine pancreatic tumor cell line) were derived from a methylcholanthrene-treated C57B1/56 mouse^56,57^. All were grown using standard cell culture methods. HPDE cells were grown in Keratinocyte media (Gibco) with 1% penicillin and streptomycin, while Panc10.05 and AsPC-1 cells were grown in RPMI 1640, L-Glutamine media (Gibco) supplemented with 1% penicillin and streptomycin, sodium pyruvate (Gibco), non-essential amino acids (Gibco), 10% FBS (ATLAS Bio), and 0.2% insulin. In accordance with NIH guidelines, cell lines were authenticated using short tandem repeat profiling at the genetic recourses core facility at Johns Hopkins University.

#### Engineered cell lines

Both lentiviral knockdown and adenoviral overexpression cell lines were generated in Panc10.05, and in some cases AsPC-1 and HDPE parental strains. For lentiviral knockdown, the hairpins used (Sigma Mission shRNA) were selected after having analyzed a minimum of three shRNAs for each gene:

shCTL NT control: 5'-CAACAAGATGAAGAGCACCAA-3'

shIIA: 5'-CCGG-GCCAAGCTCAAGAACAAGCAT-CTCGAG-ATGCTTGTTCTTGAGCTTGGC-TTTTT-3'

shIIC: 5'-CCGG-GCTCAAATATGAGGCCACAAT-CTCGAG-ATTGTGGCCTCATATTTGAGCTTTTTTG-
3'

shACTN4:5'-CCGG-CAGGACATGTTCATCGTCCAT-CTCGAG-ATGGACGATGAACATGTCCTG-TTTTTG-3'

Target plasmids were co-transfected with generation 2.0 lentiviral packaging plasmids psPAX.2 and pMD2.G via Transit 20/20 (Mirrus) transfection reagent into Lenti-X HEK293t cells. 16 hrs after transfection, the media was changed to fresh DMEM (10% FBS/1% penicillin-streptomycin). Virus-containing media was harvested after an additional 24 hrs for lentiviral infection to target cells. Positively infected cells were then selected for with 1 or 5 ng/ml puromycin in Panc10.05 or AsPC-1 cells, respectively, for 5 days as determined by kill-curve analysis. Knockdown was confirmed with western analysis.

For overexpression using the adenoviral system, fluorescent adenovirus for the expression of GFP-MYH9, GFP-MYH10, MYH14-GFP, mCherry-ACTN1, GFP-ACTN4, and GFP control were purchased from Vector BioLabs, Malvern, PA. Fluorescent filamin constructs were not generated due to their large size. Optimal multiplicity of infection (MOI= # of virus particles/cell) was first calculated by plating equal numbers of cells in a 96-well plate, then titrating virus between 0 and 200 MOi and observing fluorescence and cell death at 48 hours. For the myosins, the optimal MOI was found to be 50, where cell death was not seen and the percent of fluorescent cells was highest. While an MOI of 50 showed the highest expression and no death for the actinin constructs, the amount of protein expressed in cells was extremely high by western analysis, and so the MOI was lowered to 10. For all studies, an MOI of 50 was used for the GFP control.

### Immunohistochemistry of patient-derived samples

#### Antibodies and reagents

Antibodies used include: myosin IIA Poly19098 (BioLegend, 909801), myosin IIB (D8H8) XP (Cell Signaling Technology, #8824), myosin IIC (D4A7) (Cell Signaling Technology, #8189), α-actinin 1 (OTI7A4) (OriGene, TA500072), α-actinin 4 (G-4) (Santa Cruz Biotechnology, sc-390205), filamin A (Cell Signaling Technology, #4762), filamin B antibody [N1] (GeneTex, GTX101206), and β-actin (8H10D10) (Cell Signaling Technology, #3700).

#### Tissue preparation

The human tissue was collected and evaluated under JHH IRB #NA_00001584. Human pancreatic cancer samples were fixed in formalin, paraffin embedded, and processed for routine histology. Additional 5μm sections were cut onto plus slides and baked prior to IHC staining.

#### Immunohistochemistry staining

Immunohistochemistry was performed as previously described ^30^. In short, warmed slides were deparaffinized in sequential zylene washes, followed by 100%, 95%, and 70% ethanol washes. Slides were incubated in 0.3% H_2_O_2_ in MeOH for 20 min, then washed twice in water. Slides are steamed in citrate buffer (pH 6.0) for 35 min. Cooled slides were washed twice in water and three times in TBST (50 mM Tris-Cl, pH 7.5; 150 mM NaCl; 0.1% Tween-20). Slides were edge dried with a Kimwipe and Serum Free Protein Block Dako X0909 (Agilent, Santa Clara, CA) was applied for 10 min. Dried slides were incubated with primary antibody for 1 hr at 22^o^C, washed in TBST three times, air-dried, and incubated in secondary antibody (Dako, K400111-2, EnVision+, HRP. Mouse or K401111 -2, En Vision+ HRP. Rabbit 1100 tests (Dako) for 20 min at 22^o^C. Slides were washed three times in TBST, followed by incubation for 2-3 min in DAB+. Stained slides were washed with water, incubated for 15 sec in hematoxylin (Sigma), and washed first with water, then ethanol, and water again, incubated in acid alcohol, with a final water rinse. Slides are incubated in bluing water, washed, and dehydrated in 70% ethanol for 2 min, 100% ethanol for 2 min, and xylenes for 1 min.

#### Scoring and imaging

Slides were scanned using the Hamamatsu Digital Scanner and white balanced in Adobe Photoshop (Adobe Systems, Inc). Samples were scored based on both intensity of staining and surface area of duct covered by staining. Each slide was visualized in its entirety to determine uniformity of staining. Five cancerous ducts and five normal ducts (when present) were selected at random. Ducts with no staining were given a 0/0, ducts with intense staining on over 50% of the ductal surface were given a 2/2, with intermediate staining lying between these two endpoints. Ducts with intense staining on less than 50% of their area were scored as 2/1, ducts with moderate staining on over 50% of their ductal surface were scored as 1/2, and those with moderate staining on less than 50% of their ductal surface were scored as 1/1. To quantify the distribution of staining across all patient samples (Fig. 2B), individual duct scores were reassigned as follows: 0/0 as a 1, 1/1 as a 2, 1/2 as a 3, 2/1 as a 4, and 2/2 as a 5. Slides were imaged on NDP.View2 NanoZoomer Digital Pathology (NDP.View2, Hamamatsu Photonics, Japan).

### Quantification of cellular myosin II paralog concentrations by western analysis

#### Myosin II tail fragment protein purification

Bacterial expression plasmids coding for an N-terminal 6xHis tag, fused to the mCherry fluorophore, fused to the assembly domains of human myosin-IIA (residues 1722-1960), human myosin-IIB (residues 1729-1976), and mouse myosin-IIC (residues 1782-2033) were generated in pBiEx1 using standard cloning techniques. Proteins were expressed in BL-21 Star^™^ (DE3) (Invitrogen) *E. coli* in LB shaking culture overnight at room temperature. Bacteria were harvested by centrifugation and lysed by lysozyme treatment followed by sonication, and the lysate was clarified by centrifugation. Polyethyleneimine (PEI) was added to a final concentration of 0.1% to precipitate nucleic acids, which were then removed by centrifugation. The myosin-II constructs were precipitated by adding ammonium sulfate to 50% saturation and centrifuging. The pellet was resuspended in column running buffer, dialyzed against the same for a minimum of 4 hours, clarified by centrifugation and filtration, and run on a Ni-NTA metal affinity column, followed by a sizing column. The constructs were then concentrated and further purified by dialyzing against assembly buffer (10 mM HEPES, pH 7.1, 50 mM NaCl) until precipitate formed, followed by centrifugation and resuspension of the pellet in storage buffer (10 mM HEPES, pH 7.1, 500 mM NaCl). Protein purity was verified by SDS-PAGE followed by Coomassie Blue staining, and concentration was quantified by UV absorbance using the calculated extinction coefficient for each protein’s amino acid sequence.

#### Quantitative western analysis

Cells were trypsinized and counted, then centrifuged into pellets containing 5x10^5^ cells each. These pellets were washed in PBS and recentrifuged, then lysed in 75 μL RIPA lysis buffer plus 15 μL 6xSDS buffer. Due to cell volume and residual PBS, the total lysate volume reached 100 μL. 10 μL of lysate was added to each well of a 7% SDS-PAGE gel, or the equivalent of 5x10^4^ cells/well. In addition, each well was spiked with a known quantity of purified myosin II tail fragment, containing the epitope region for the antibodies used, with sequential 2-fold dilutions. A 7% gel was used because it allowed for optimal transfer of both the large molecular weight endogenous myosin II and the smaller molecular weight purified tail fragment out of the gel. Transfer was most effective at a constant 45V for 16 hrs, using PVDF membranes to prevent smaller protein pass-through and verifying complete transfer of larger proteins by performing a Coomassie stain to verify that no protein was left in the gel following transfer. The average volume of an individual cell for each cell type was determined from the micropipette aspiration images, where cell radius is measured, and assuming the cell shape to be a sphere prior to aspiration. For each experiment, a standard curve was created from the spiked tail fragment to determine the total number of moles of endogenous myosin II in each lane. The number of cells per lane multiplied by the average volume of a single cell gave the total cell volume per lane, and concentration was determined as a ratio of these two values. Antibodies used were the same as those for immunohistochemistry (described above).

### Micropipette aspiration assay for mechanoresponse and mechanics measurements and realtime deformability cytometry measurements

The instrumental and experimental setups have been described previously^42,58^. MPA assays and RT-DC measurements were all carried out in growth media for cortical tension measurements or Leibovitz L-15 media (Gibco) when fluorescence was quantified.

#### Measurements of mechanosensory accumulation of proteins

A pressure difference was generated by adjusting the height of a motor-driven water manometer. Mammalian cells expressing desired fluorescent proteins were loaded into the observation chamber, which was filled with Leibovitz L-15 Medium w/o phenol red (Gibco). Cells were deformed using a pressure of 0.3 nN/μm^2^ and recorded for 5 min. Pressures higher than this often led to blebbing or the separation of cell membrane from the cortex. All cells which demonstrated blebbing during recording were discarded. Images were collected with an Olympus IX81 microscope equipped with MetaMorph software and analyzed using ImageJ (National Institutes of Health). After background correction, the fluorescence intensity at the accumulation site inside the micropipette was normalized against the opposite cortex of the cell (I_p_/I_o_). The peak I_p_/I_o_ value during the 5 min timecourse was then normalized to the I_p_/I_o_ value at t=0 to adjust for initial variations in cortical fluorescence (Normalized I_p_/I_o_).

#### Cortical tension measurements

Pressure was applied to the cell cortex with a micropipette (6‐ to 8-μm radius; R_p_) to the equilibrium pressure (ΔP), where the length of the cell inside the pipette (L_p_) was equal to R_p_. The effective cortical tension (T_eff_) was calculated by applying the Young–Laplace equation: ΔP = 2T_eff_ (1/R_p_ – 1/R_c_), where R_c_ is the radius of the cell and ΔP is the equilibrium pressure when L_p_ = R_p_^58^ Images were collected with an Olympus IX81 microscope equipped with MetaMorph software and analyzed using ImageJ (rsb.info.nih.gov/ij).

#### Real-time deformability cytometry

Mechanical measurements of thousands of cells were obtained as previously described ^42^ Approximately 10^6^ cells were trypsinized, spun, resuspended in media, and incubated at 37μC for 10 min prior to loading onto the AcCellerator (Zellmechanik Dresden), using a 30μm channel. Deformation and cell size data was collected in real-time at three different flow-rates and analyzed using ShapeOut (Zellmechanik Dresden; available at https://github.com/zellmechanik-dresden/ShapeOut). Differences in deformation were plotted as a probability distribution in R (r-project.org/), and the elastic modulus, based on the median of the deformation and area populations, the channel width, viscosity, and flow rate was calculated ^43^

### Imaging and Image Analysis

Imaging was performed in culture media or Leibovitz L-15 media without phenol red (Gibco) and 10% FBS for mechanoresponse and lattice light sheet experiments. Confocal imaging was performed on a Zeiss 510 Meta microscope with a 63X (1.4 N.A.) objective (Carl Zeiss). Epifluorescence imaging was performed with an Olympus IX81 microscope using a 40X (1.3 N.A.) objective and a 1.6X optovar (Olympus), as previously described. Image analysis was performed with ImageJ (rsb.info.nih.gov/ij). Datasets were independently analyzed by multiple investigators.

### Single cell assays

#### 2D random migration

AsPC-1 cells were plated at a sub-confluent concentration in a 24-well tissue culture plate (<5,000 cells) and incubated overnight in growth media (see above). Prior to imaging, cells media was changed to Leibovitz L-15 media without phenol red (Gibco) containing 10% FBS and 1% penicillin/streptomycin. Cells were imaged using Molecular Devices IXM High Content Imager 10X objective (NA) every 30 min for 24 hrs. Cell roundness, velocity and area were quantified using ImageJ.

#### Retrograde flow

AsPC-1 cells were grown on collagen-I coated (50 μg/ml) 5-mm coverslips for 16 hrs and then treated with 100-nM SiR-Actin (Cytoskeleton, Inc) and 1 μM Verapamil for 4 hrs in Leibovitz L-15 without phenol red media, with or without 4-HAP (500 nM). Coverslips were transferred to the imaging chamber of the Lattice Light-Sheet Microscope (LLSM) (Intelligent Imaging Innovations) containing fresh Leibovitz L-15 media without SiR-Actin and verapamil plus the corresponding 4-HAP concentration. Cells were imaged for 3-5 min at 2-3-sec intervals, 50-150 planes per 3D stack, with a Nikon CFI75 Apochromat 25X/1.1 water-dipping objective. Retrograde actin flow was measured using ImageJ (rsb.info.nih.gov/ij). Datasets were independently analyzed by multiple investigators.

#### Transwell assays

AsPC-1 cells were plated in 6.5-mm PET membrane transwell inserts with 8μm pores (Costar #3464) in a 24-well plate at a concentration of 5,000 cells per well. Cells were allowed to adhere overnight in AsPC-1 media. Cell media was then changed to serum-free RPMI 1640 to starve cells for 18 hrs. Following starvation, cells were stimulated by changing media in the top chamber to fresh serum-free RPMI 1640 ±500 nM 4-HAP and complete AsPC-1 media ±500 nM 4HAP. Cells were then incubated for 24 hrs at 37 degrees C/5% CO_2_ and then fixed in 4% paraformaldehyde, permeabilized in 0.1% Triton X-100 and stained with 1 μg/ml DAPI. Prior to imaging, the top chamber was swabbed with a cotton-tip swab and washed to remove cells that did not translocate. A total of five random fields per transwell insert were imaged using a 10X objective (NA) and nuclei were averaged.

### Tissue spheroid generation, staining, and quantification

#### Tissue spheroid generation

Tissue spheroids were grown by plating Panc10.05 cells on a drop of Matrigel (Becton-Dickinson) in an 8-well slide chamber (6,500 cells per well) and grown in RPMI 1640 media (2% serum, 2% matrigel, and 10 ng/ml EGF). Spheroids were grown for 14 days with regular media changes, then aspirated off the surface of the matrigel, washed with ice-cold PBS with mild centrifugation (4,000 RCF, 5 min). For all 2D spreading assays, spheroids were then plated on 50 μg/ml collagen-coated 8-well coverslips (MatTek) and incubated for 48 hrs in complete PANC media with 500 nM 4-HAP or PBS control. For 3D invasion assays, spheroids were resuspended in a 1.5 mg/ml collagen solution (Life Technologies) and then plated in 8-well chambered coverslips. Spheroids were incubated in complete PANC media with 500 nM 4-HAP or PBS control for 48 hrs. All spheroid samples were fixed in 4% paraformaldehyde for 15 min at 25°C and permeabilized in 0.1% Triton X-100 for 15 min at 25°C. The actin cytoskeleton was visualized with rhodamine-phalloidin (5 μM) for 30 min. Two-dimensional spreading was visualized with Olympus Spinning disk microscope (40X oil objective, 1.30NA) or Nikon-NSIM (100X objective, 1.4NA) with an Andor EMCCD camera controlled by NIS-Elements software. Three-dimensional invasion through collagen was visualized with an Olympus Spinning Disk microscope (20X air, 0.4NA)

#### Spheroid cortex fluorescence quantification

Quantifications of 2D spheroids plated on a thin layer of collagen was performed using a custom-designed Matlab script (Mathworks, Natick, MA). A maximum intensity projection of 20 z-slices was extracted from each of the images and converted to 16-bit grayscale before enhancing the pixel intensity (Matlab command: histeq). The enhanced image was then segmented into foreground and background using the Chan-Vese method (activecountour), followed by filtering of small regions (bwareaopen), and morphological erosion (imerode). The boundary from this object was then obtained (bwboundary) and average background intensity was subtracted. This method was used for more than 80% of the images. If this boundary did not accurate reflect the shape of the spheroid, a manual tracing option was offered. This option allowed the user to select a region of interest surrounding the outer edge of the spheroid (roipoly). Anything outside of this boundary was masked and subsequently not considered (regionfill). Following this, a similar binarization/background subtraction regime was implemented on the unmasked region and the boundary was traced.

For each point along the boundary, a line perpendicular to the edge was computed and the intensity along this line was used to linearize the cortex. The resultant matrix was converted into a grayscale image. The Hough Transform was used to determine the continuity of the actin belt along the edge of the cortex. Linearized cortex images were binarized using a threshold chosen to best differentiate belt from non-belt for each set of images (imbinarize), followed by filtering of small regions (bwareaopen), and morphological dilation (imdilate). The Hough Transform was performed on this image (hough), and Hough peaks and lines identified (houghpeaks, houghlines). These were used to create a continuity score defined by (Σ Hough line lengths)/(number of lines + length of image). In this metric, higher scores indicate a more continuous actin belt at the cortex. Additionally, we computed: a coarseness index of the cortex (std2) to describe actin distribution, the percentage of white pixels of the binarized image to characterize actin belt thickness, and the ratio of white pixels to gray pixels (greater than 0.15) to describe the distribution of fluorescence.

### Mouse studies

#### Hemi-splenectomies

Hemi-splenectomies were performed on athymic *NCr-nu/nu* mice (Charles River Laboratories) with low passage AsPC-1 cells as previously described^59^. In short, laparotomies were performed on anesthetized mice in which the upper pole of a divided spleen was reinserted into the peritoneum while 1X10^7^ AsPC-1 cells prepared in phosphate buffered saline were injected into the lower splenic pole, chased by an equal volume of phosphate buffered saline. The pancreas and splenic vessels were ligated and the peritoneum was closed.

AsPC-1 cells were pretreated with 50 μM 4-HAP or PBS 24 hr prior to injection during hemisplenectomies. Mice were also weighed and treated intraperitoneally with 200 μl of 5 mg/ml of 4-HAP or 200 μl PBS two days prior to surgery, and then treatment was continued every other day, starting on day 1 post-surgery. Mice were sacrificed upon spontaneous death of the first mouse. Samples from the mice of both this study and the survivability study were harvested as described below.

Livers were washed, weighed, photographed, and fixed in 10% formalin in PBS for 48 hrs, embedded in paraffin blocks, and sectioned as 4μm thick slides. Mice were housed and handled according to approved Institutional Animal Care and Use Committee protocols.

#### Mouse liver tumor quantification

Quantifications of mouse livers were performed using a custom-designed Matlab script. RGB images were first separated into their three component channels. The imageSegmenter tool on Matlab was used to segment the individual channels identifying the area of the whole liver, the area of the tumor, and the area/location of the glare on the images, which was removed from the tumor area. The percent coverage of tumor was calculated.

## Acknowledgments

We thank Drs. Hoku West-Foyle of the JHMI Microscope Facility and Marc Edwards for assistance in image acquisition, Drs. Martin Kräter, Christoph Herold, and Angela Jacobi of TU Dresden for tissue culture maintenance and assistance with RT-DC, Dr. Hoku West-Foyle again for reagents (mammalian myosin II tail fragments), members of the Jaffee lab for helpful conversations on mouse *in vivo* data analysis, and members of the Robinson lab for helpful conversations over the course of this project, especially Jennifer Nguyen for help with R. This work was supported by the NIH (GM66817, GM109863), the Sol Goldman Foundation, a Johns Hopkins Discovery Grant, and the Alexander-von-Humboldt Foundation (AvH Professorship to JG).

## References

1 Remmerbach, T. W. et al. Oral cancer diagnosis by mechanical phenotyping. Cancer Res 69, 1728–1732, doi:10.1158/0008-5472.CAN-08-4073 (2009).

2 Guck, J. et al. Optical deformability as an inherent cell marker for testing malignant transformation and metastatic competence. Biophys J 88, 3689–3698, doi:10.1529/biophysj.104.045476 (2005).

3 Cross, S. E., Jin, Y. S., Rao, J. & Gimzewski, J. K. Nanomechanical analysis of cells from cancer patients. Nat Nanotechnol 2, 780–783, doi:10.1038/nnano.2007.388nnano.2007.388 [pii] (2007).

4 Staunton, J. R., Doss, B. L., Lindsay, S. & Ros, R. Correlating confocal microscopy and atomic force indentation reveals metastatic cancer cells stiffen during invasion into collagen I matrices. Sci Rep 6, 19686, doi:10.1038/srep19686 (2016).

5 Suresh, S. Biomechanics and biophysics of cancer cells. Acta Biomater. 3, 413–438 (2007).

6 Kim, T. H. et al. Cancer cells become less deformable and more invasive with activation of beta-adrenergic signaling. J Cell Sci 129, 4563–4575, doi:10.1242/jcs.194803 (2016).

7 Nguyen, A. V. et al. Stiffness of pancreatic cancer cells is associated with increased invasive potential. Integr Biol (Camb) 8, 1232–1245, doi:10.1039/c6ib00135a (2016).

8 Lu, P., Weaver, V. M. & Werb, Z. The extracellular matrix: a dynamic niche in cancer progression. J Cell Biol 196, 395–406, doi:10.1083/jcb.201102147 (2012).

9 Jones, S. et al. Core signaling pathways in human pancreatic cancers revealed by global genomic analyses. Science 321, 1801–1806, doi:10.1126/science.11643681164368 [pii] (2008).

10 Maimaiti, Y. et al. Overexpression of cofilin correlates with poor survival in breast cancer: A tissue microarray analysis. Oncol Lett 14, 2288–2294, doi:10.3892/ol.2017.6413 (2017).

11 Pappa, K. I. et al. Proteomic Analysis of Normal and Cancer Cervical Cell Lines Reveals Deregulation of Cytoskeleton-associated Proteins. Cancer Genomics Proteomics 14, 253–266, doi:10.21873/cgp.20036 (2017).

12 Wang, W. S. et al. The expression of CFL1 and N-WASP in esophageal squamous cell carcinoma and its correlation with clinicopathological features. Dis Esophagus 23, 512–521, doi:10.1111/j.1442-2050.2009.01035.x (2010).

13 Kikuchi, S. et al. Expression and gene amplification of actinin-4 in invasive ductal carcinoma of the pancreas. Clin Cancer Res 14, 5348–5356, doi:10.1158/1078-0432.CCR-08-0075 (2008).

14 Fukumoto, M., Kurisu, S., Yamada, T. & Takenawa, T. alpha-Actinin-4 enhances colorectal cancer cell invasion by suppressing focal adhesion maturation. PLoS One 10, e0120616, doi:10.1371/journal.pone.0120616 (2015).

15 Honda, K. et al. Actinin-4 increases cell motility and promotes lymph node metastasis of colorectal cancer. Gastroenterology 128, 51–62 (2005).

16 Liu, X. & Chu, K. M. alpha-Actinin-4 promotes metastasis in gastric cancer. Lab Invest 97, 1084–1094, doi:10.1038/labinvest.2017.28 (2017).

17 Shiraishi, H. et al. Actinin-4 protein overexpression as a predictive biomarker in adjuvant chemotherapy for resected lung adenocarcinoma. Biomark Med, doi:10.2217/bmm-2017-0150 (2017).

18 Gao, Y. et al. ACTN4 and the pathways associated with cell motility and adhesion contribute to the process of lung cancer metastasis to the brain. BMC Cancer 15, 277, doi:10.1186/s12885-015-1295-9 (2015).

19 Yamaguchi, H. et al. Actinin-1 and actinin-4 play essential but distinct roles in invadopodia formation by carcinoma cells. Eur J Cell Biol, doi:10.1016/j.ejcb.2017.07.005 (2017).

20 Iguchi, Y. et al. Filamin B Enhances the Invasiveness of Cancer Cells into 3D Collagen Matrices. Cell Struct Funct 40, 61–67, doi:10.1247/csf.15001 (2015).

21 Steinhardt, A. A. et al. Expression of Yes-associated protein in common solid tumors. Hum Pathol 39, 1582–1589, doi:10.1016/j.humpath.2008.04.012S0046-8177(08)00194-9 [pii] (2008).

22 Bai, H. et al. Yes-Associated Protein impacts adherens junction assembly through regulating actin cytoskeleton organization. Am. J. Physiol. – Gastrointest. Liver Phys. 311, G369–G411 (2016).

23 Maitra, A. et al. Multicomponent analysis of the pancreatic adenocarcinoma progression model using a pancreatic intraepithelial neoplasia tissue microarray. Mod Pathol 16, 902–912, doi:10.1097/01.MP.0000086072.56290.FB (2003).

24 Sun, Q. et al. Competition between human cells by entosis. Cell Res. 24, 1299–1310 (2014).

25 Tang, Y. et al. 14-3-3zeta up-regulates hypoxia-inducible factor-1alpha in hepatocellular carcinoma via activation of PI3K/Akt/NF-small ka, CyrillicB signal transduction pathway. Int J Clin Exp Pathol 8, 15845–15853 (2015).

26 Su, C. H. et al. 14-3-3sigma exerts tumor-suppressor activity mediated by regulation of COP1 stability. Cancer Res 71, 884–894, doi:10.1158/0008-5472.CAN-10-2518 (2011).

27 Tong, S. et al. 14-3-3zeta promotes lung cancer cell invasion by increasing the Snail protein expression through atypical protein kinase C (aPKC)/NF-kappaB signaling. Exp Cell Res 348, 1-9, doi:10.1016/j.yexcr.2016.08.014 (2016).

28 Yang, X. et al. 14-3-3zeta positive expression is associated with a poor prognosis in patients with glioblastoma. Neurosurgery 68, 932–938, doi:10.1227/NEU.0b013e3182098c30 (2011).

29 Zhou, Q. et al. 14-3-3 coordinates microtubules, Rac and myosin II to control cell mechanics and cytokinesis. Curr. Biol. 20, 1881–1889, doi:10.1016/j.cub.2010.09.048 (2010).

30 Jhaveri, D. T. et al. Using Quantitative Seroproteomics to Identify Antibody Biomarkers in Pancreatic Cancer. Cancer Immunol Res 4, 225–233, doi:10.1158/2326-6066.CIR-15-0200-T (2016).

31 Luo, T., Mohan, K., Iglesias, P. A. & Robinson, D. N. Molecular mechanisms of cellular mechanosensing. Nat Mater 12, 1064–1071, doi:10.1038/nmat3772 (2013).

32 Surcel, A. et al. Pharmacological activation of myosin II paralogs to correct cell mechanics defects. Proc Natl Acad Sci U S A 112, 1428–1433 (2015).

33 Effler, J. C. et al. Mitosis-specific mechanosensing and contractile-protein redistribution control cell shape. Curr Biol 16, 1962–1967, doi:10.1016/j.cub.2006.08.027 (2006).

34 Kee, Y. S. et al. A mechanosensory system governs myosin II accumulation in dividing cells. Mol Biol Cell 23, 1510–1523 (2012).

35 Luo, T. et al. Understanding the cooperative interaction between myosin II and actin cross-linkers mediated by actin filaments during mechanosensation. Biophys. J. 102, 238–247 (2012).

36 Schiffhauer, E. S. et al. Mechanoaccumulative Elements of the Mammalian Actin Cytoskeleton. Curr Biol 26, 1473–1479, doi:10.1016/j.cub.2016.04.007 (2016).

37 Thomas, D. G. & Robinson, D. N. The fifth sense: Mechanosensory regulation of alpha-actinin-4 and its relevance for cancer metastasis. Semin Cell Dev Biol, In press, doi:10.1016/j.semcdb.2017.05.024 (2017).

38 Pei, H. et al. FKBP51 affects cancer cell response to chemotherapy by negatively regulating Akt. Cancer Cell 16, 259–266, doi:10.1016/j.ccr.2009.07.016 (2009).

39 Uhlen, M. et al. A pathology atlas of the human cancer transcriptome. Science 357, doi:10.1126/science.aan2507 (2017).

40 Wu, J.-Q. & Pollard, T. D. Counting cytokinesis proteins globally and locally in fission yeast. Science 310, 310–314 (2005).

41 Robinson, D. N., Cavet, G., Warrick, H. M. & Spudich, J. A. Quantitation of the distribution and flux of myosin-II during cytokinesis. BMC Cell Biol 3, 4 (2002).

42 Otto, O. et al. Real-time deformability cytometry: on-the-fly cell mechanical phenotyping. Nat Methods 12, 199–202, doi:10.1038/nmeth.3281 (2015).

43 Mokbel, M. et al. Numerical Simulation of Real-Time Deformability Cytometry To Extract Cell Mechanical Properties. ACS Biomater. Sci. Eng., doi:10.1021/acsbiomaterials.6b00558 (2017).

44 Reichl, E. M. et al. Interactions between myosin and actin crosslinkers control cytokinesis contractility dynamics and mechanics. Curr Biol 18, 471–480, doi:10.1016/j.cub.2008.02.056 (2008).

45 Liu, T. et al. Downregulation of nonmuscle myosin IIA expression inhibits migration and invasion of gastric cancer cells via the cJun Nterminal kinase signaling pathway. Mol Med Rep 13, 1639–1644, doi:10.3892/mmr.2015.4742 (2016).

46 Schramek, D. et al. Direct in vivo RNAi screen unveils myosin IIa as a tumor suppressor of squamous cell carcinomas. Science 343, 309–313, doi:10.1126/science.1248627 (2014).

47 Conti, M. A. et al. Conditional deletion of nonmuscle myosin II-A in mouse tongue epithelium results in squamous cell carcinoma. Sci Rep 5, 14068, doi:10.1038/srep14068 (2015).

48 Huelsmann, S., Rintanen, N., Sethi, R., Brown, N. H. & Ylanne, J. Evidence for the mechanosensor function of filamin in tissue development. Sci Rep 6, 32798, doi:10.1038/srep32798 (2016).

49 Zhou, X. et al. Filamin B deficiency in mice results in skeletal malformations and impaired microvascular development. Proc Natl Acad Sci U S A 104, 3919–3924, doi:10.1073/pnas.0608360104 (2007).

50 Holterhoff, C. K., Saunders, R. H., Brito, E. E. & Wagner, D. S. Sequence and expression of the zebrafish alpha-actinin gene family reveals conservation and diversification among vertebrates. Dev Dyn 238, 2936–2947, doi:10.1002/dvdy.22123 (2009).

51 Wylie, S. R. & Chantler, P. D. Myosin IIC: a third molecular motor driving neuronal dynamics. Mol Biol Cell 19, 3956–3968, doi:10.1091/mbc.E07-08-0744 (2008).

52 Recuenco, M. C. et al. Nonmuscle Myosin II Regulates the Morphogenesis of Metanephric Mesenchyme-Derived Immature Nephrons. J Am Soc Nephrol 26, 1081–1091, doi:10.1681/ASN.2014030281 (2015).

53 Lalwani, A. K. et al. Localization in stereocilia, plasma membrane, and mitochondria suggests diverse roles for NMHC-IIa within cochlear hair cells. Brain Res 1197, 13–22, doi:10.1016/j.brainres.2007.12.058 (2008).

54 Reichl, E. M. et al. Interactions between myosin and actin crosslinkers control cytokinesis contractility dynamics and mechanics. Curr. Biol. 18, 471–480 (2008).

55 Srivastava, V., Iglesias, P. A. & Robinson, D. N. Cytokinesis: Robust cell shape regulation. Semin Cell Dev Biol 53, 39–44, doi:10.1016/j.semcdb.2015.10.023 (2016).

56 Corbett, T. H. et al. Induction and chemotherapeutic response of two transplantable ductal adenocarcinomas of the pancreas in C57BL/6 mice. Cancer Res 44, 717–726 (1984).

57 Schmidt, T. et al. Intratumoral immunization with tumor RNA-pulsed dendritic cells confers antitumor immunity in a C57BL/6 pancreatic murine tumor model. Cancer Res 63, 8962–8967 (2003).

58 Kee, Y.-S. & Robinson, D. N. Micropipette Aspiration for Studying Cellular Mechanosensory Responses and Mechanics. Dictyostelium Protocols II: Methods Mol. Biol. 983, 367–382 (2013).

59 Soares, K. C. et al. A preclinical murine model of hepatic metastases. J Vis Exp, 51677, doi:10.3791/51677 (2014).

